# Cell-based assays and comparative genomics revealed the conserved and hidden effects of *Wolbachia* on insect sex determination

**DOI:** 10.1101/2024.02.12.579973

**Authors:** Hiroshi Arai, Benjamin Herran, Takafumi N. Sugimoto, Mai Miyata, Tetsuhiko Sasaki, Daisuke Kageyama

## Abstract

It is advantageous for maternally transmitted endosymbionts to skew the sex ratio of their hosts toward females. Some endosymbiotic bacteria such as *Wolbachia* cause their insect hosts to exclusively produce female offspring through male killing or feminization. In some lepidopteran insects, male killing is achieved by affecting the sex-determining process in males, and a unique mechanism of male killing and its functional link with feminization have been implicated. However, comparative analysis of these phenotypes is often difficult because they have been analyzed in different host–symbiont systems and transinfection of *Wolbachia* across different hosts is often challenging. In this study, we demonstrated the effects of nine *Wolbachia* strains on the splicing of sex-determining genes in Lepidoptera by fixing the host genetic background using a cell culture system. Cell transinfection assays confirmed that three male killing-inducing *Wolbachia* strains and one feminization-inducing *Wolbachia* strain increased the female-type splicing products of the core sex-determining genes *doublesex*, *masculinizer*, and *zinc finger protein 2*. Regarding *Wolbachia* strains that do not induce male killing/feminization, three had no effect on these sex-determining genes, whereas two strains induced female-type splicing of *masculinizer* and *doublesex* but not *zinc finger protein 2*. Comparative genomics confirmed that homologs of *oscar*, the *Wolbachia* gene responsible for male killing in *Ostrinia*, were encoded by male killing/feminizing *Wolbachia* strains, but not by non-male killing/non-feminizing strains. These results demonstrated the conserved effects underlying male killing and feminization induced by *oscar*-bearing *Wolbachia*, and suggested other potential mechanisms that *Wolbachia* might employ to manipulate host sex.

**IMPORTANCE:** Arthropods commonly carry maternally transmitted microbial symbionts such as *Wolbachia*. The lack of paternal transmission frequently led to the evolution of reproductive parasitism traits, namely the manipulation of host reproduction in favor of female hosts, substantiated by male killing and feminization. Although *Wolbachia* induces these phenotypes in a wide range of insects, the underlying mechanisms, diversity, and commonality remain largely unclear. In this study, we used a combination of transinfection assays and comparative genomics to reveal the conserved effects of male killing and feminizing *Wolbachia* strains on lepidopteran sex determination. Furthermore, we demonstrated that some non-male killing/non-feminizing *Wolbachia* strains also have an inherent ability to influence sex determination, albeit in a different manner, suggesting the potential for multiple mechanisms to manipulate host sex. This study also implied the frequent evolution of host suppressors against *Wolbachia*-induced reproductive manipulation.

## INTRODUCTION

Most animal species have distinct male and female forms that are determined by their genomic information during early development in a process called sex determination (1–2). However, in many cases, sex-determining systems are manipulated or overridden by maternally transmitted cytoplasmic elements (3–6). In arthropods, several bacterial endosymbionts manipulate host reproduction by killing (i.e., male killing [MK]) or feminizing males (feminization) (7–11). The maternally inherited bacterium *Wolbachia* (Alphaproteobacteria) is a genus of typical microbes that manipulate host reproduction through various mechanisms (12). For example, *Wolbachia* species induce MK and FM in diverse insects belonging to phylogenetically distinct taxa (e.g., Diptera, Lepidoptera, and Coleoptera) and other arthropods (8-9; 13-16). *Wolbachia* species are estimated to inhabit at least 40% of all insect species, making them among the most widespread endosymbionts globally (12, 17).

It has been hypothesized that the mechanisms underlying MK and FM are associated with sex determination cascades in insects (18). Indeed, some MK *Wolbachia* strains directly interact with sex determination systems in certain insects (19–22). In *Ostrinia* and *Homona* male moths, MK *Wolbachia* strains affect the splicing of *doublesex* (*dsx*), a downstream master regulator of sex determination and differentiation, resulting in the production of the ‘female-type’ isoform *dsxF* (19–22). In *Ostrinia* males, an MK *Wolbachia* strain (*w*Sca) also induces the female-specific transcript variant of the autosomal CCCH-type zinc finger motif-encoding gene *zinc finger protein 2* (*znf2*) (23), which is an upstream factor of *dsx* in *Bombyx mori* (24). Similarly, the FM *Wolbachia* strain *w*Fem induces *dsxF* production in *Eurema* butterflies (25). The ‘mismatch’ between genetic sex (male: ZZ sex chromosome constitution) and phenotypic sex (female: based on *dsxF*) leads to the collapse of dosage compensation, a system that equalizes the sex-linked (Z chromosome-linked) gene dose between males and females (20–21). In *Ostrinia*, these changes are caused by the *Wolbachia* protein Oscar, which interacts with and degrades the Z-linked CCCH-type zinc finger motif-encoding gene *masculinize* (*masc*). This gene regulates both dosage compensation and *dsx* splicing patterns in Lepidoptera (26–27). Furthermore, an *oscar* homolog was found in the prophage region of an MK *Wolbachia* strain in *Homona* moths (28). These findings imply a general role for Oscar in MK in lepidopteran insects, which has not been demonstrated. Based on the *Wolbachia* strains sequenced to date, *oscar* is not widely conserved among MK *Wolbachia* strains (27–28). In addition, transgenically overexpressed Oscar does not function in *Drosophila* flies, which do not employ *masc* in sex determination. Instead, another *Wolbachia* factor, Wmk, induces male lethality (28–29). In contrast to Oscar, Wmk homologs are widely conserved among *Wolbachia* strains regardless of their phenotypes (28–29). These findings imply that *Wolbachia* uses multiple mechanisms to induce MK in insects, but their similarity/diversity and functional link with FM remain elusive.

The induction of *Wolbachia*-induced reproductive manipulation is deeply associated with the host genetic background (30–33). MK and FM can significantly distort the sex ratio of the host population toward females (13, 31, 34–36). Under extremely female-biased sex ratios, male production is strongly selected, and the development of host resistance to endosymbiont-induced MK has been observed in some insects, such as butterflies, ladybugs, lacewings, and planthoppers (36–40). However, the subsequent transfer of the ‘suppressed’ male killer into the genetic background of a new host resulted in the reemergence of MK (30). Therefore, the absence of obvious *Wolbachia*-induced phenotypes in the natural host does not necessarily indicate the absence of *Wolbachia* MK capabilities. Thus, investigations of the inherent MK/FM abilities of *Wolbachia* strains are warranted to elucidate the genetic basis of these phenotypes.

In this study, we revealed a conserved and hidden feminizing effect induced by *Wolbachia* strains in Lepidoptera. Transinfection assays using a recently established cell culture system (23) revealed that MK/FM *Wolbachia* strains altered the splicing patterns of genes involved in the moth sex determination cascade. Genomic analyses confirmed the presence of Oscar in MK/FM *Wolbachia* strains. Furthermore, an inherent feminizing ability was observed in Oscar-deficient *Wolbachia* strains that do not induce MK or FM in their native hosts. Based on these findings, we discussed the underlying mechanisms of MK and FM in Lepidoptera and highlighted the diversity of causative factors.

## RESULTS

### Transinfection assays revealed the feminizing abilities of *Wolbachia* strains in *Ostrinia* male cells

The OsM1 cell line derived from a male moth (*Ostrinia scapulalis*) (23) was transinfected with the native *Wolbachia* strain *w*Sca and eight non-native *Wolbachia* strains from lepidopteran insects (**Table 1**). Consistent with previous findings (23), transinfection with *w*Sca (native male killer) induced female-type splicing patterns in *dsx* (*OsdsxF*), *masc* (*OsmascF*), and *znf2* (*Osznf2F*) at 6 weeks post-transinfection (6 wpt) compared to the findings in non-transinfected controls (Steel– Dwass test; *Osdsx*: *p* = 0.02; *Osmasc*: *p* = 0.02; *Osznf2*: *p* = 0.02; **Figure 1**). Albeit less or more conspicuously, *w*CauA, *w*Hm-t, and *w*Fem, having MK/FM abilities in native or non-native hosts, also increased the expression of *OsdsxF* (*w*CauA: *p* = 0.01, *w*Hm-t: *p* = 0.04, *w*Fem: *p* = 0.02), *OsmascF* (*w*CauA: *p* = 0.01, *w*Hm-t: *p* = 0.04, *w*Fem: *p* = 0.02), and *Osznf2F* (*w*CauA: *p* = 0.01, *w*Hm-t: *p* = 0.04, *w*Fem: *p* = 0.02) at 6 wpt compared to the control levels. Cells transinfected with either *w*Kue, *w*CauB, or *w*CI (inducing cytoplasmic incompatibility but not MK/FM in native hosts) continued to exhibit male-type splicing patterns for *Osdsx* (*w*Kue: *p* = 0.48, *w*CauB: *p* = 0.99, *w*CI: *p* = 0.99), *Osmasc* (*w*Kue: *p* = 0.99, *w*CauB: *p* = 0.99, *w*CI: *p* = 0.44), and *Osznf2* (*w*Kue: *p* = 0.07, *w*CauB: *p* = 0.13, *w*CI: *p* = 0.80) similarly as observed in *Wolbachia*-free control cells. Unexpectedly, *w*Hm-c, which does not induce MK/FM in its host (41–42), upregulated *OsdsxF* (*p* = 0.02) and *OsmascF* (*p* = 0.02), but not *Osznf2F* (*p* = 0.07), at 6 wpt compared to the control levels.

**Fig. 1.**
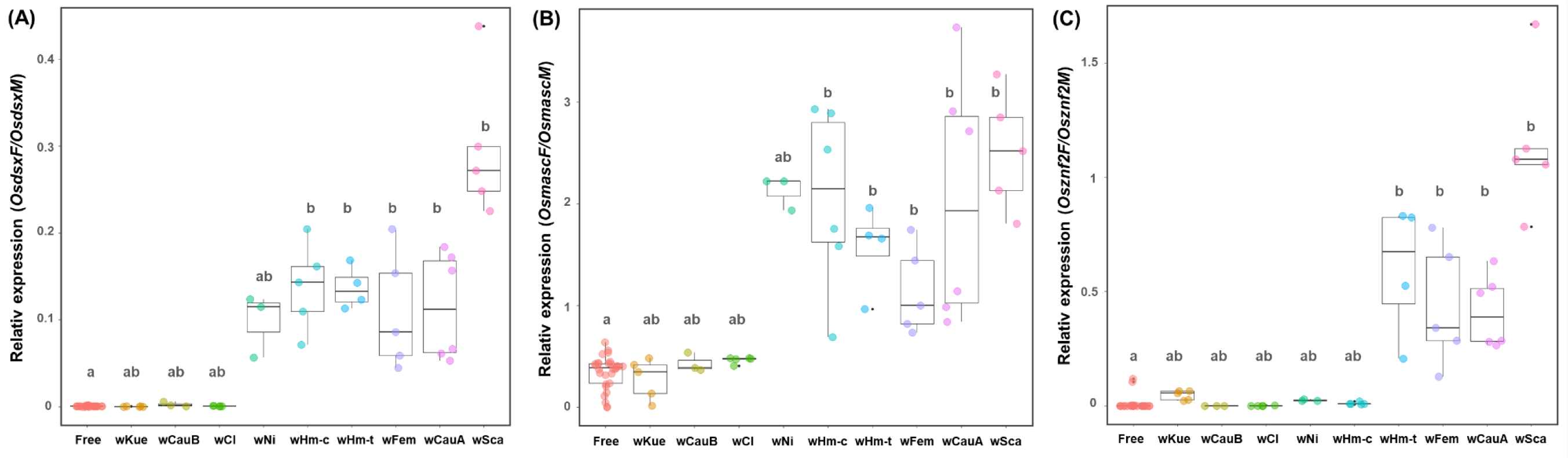
Feminizing effects of *Wolbachia* strains on *Ostrinia* male cells Relative expression of *OsdsxF/OsdsxM* (A), *OsmascF/OsmascM* (B), and *Osznf2F/Osznf2M* (C) was quantified in *Wolbachia*-transinfected and control cells. Different letters above the whisker plots indicate significant differences based on the Steel–Dwass test (*p* < 0.05). Dot plots indicate replicates.

**Table 1.**
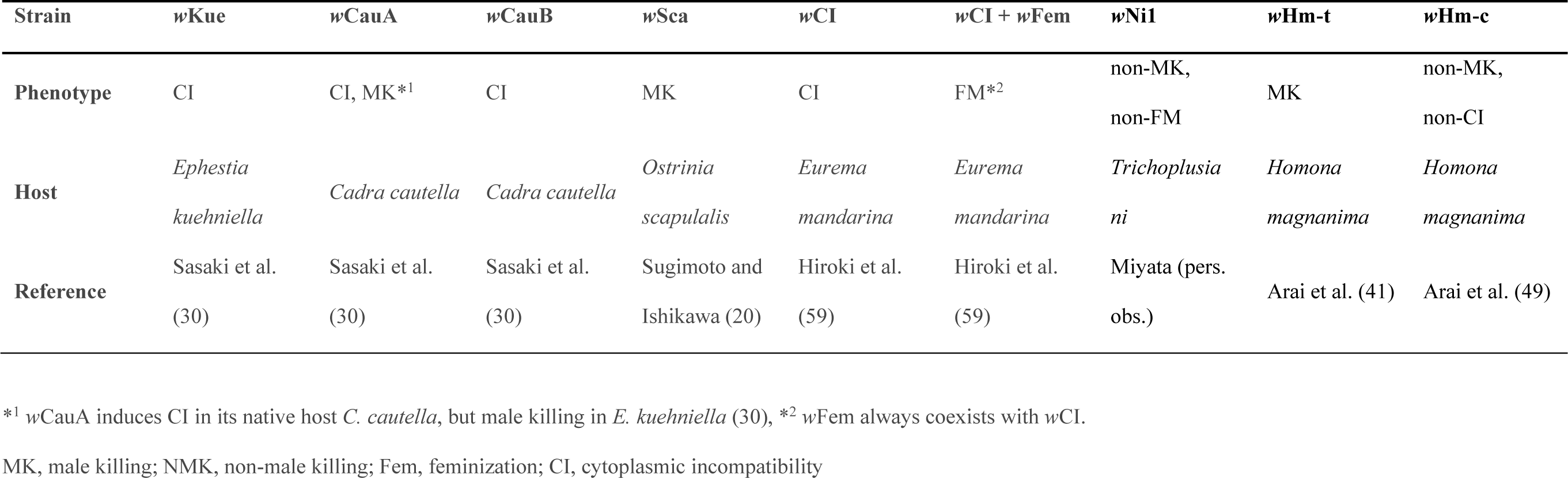
Characteristics of the *Wolbachia* strains used in this study.

Similar patterns were also observed for the non-MK/FM *w*Ni1 strain in *Trichoplusia ni*, including the upregulation of *OsdsxF* (*p* = 0.13) and *OsmascF* (*p* = 0.13), albeit without significance, whereas no effect was observed on *Osznf2F* (*p* = 0.31; **Figure 1**).

### Oscar is conserved among MK and FM *Wolbachia* strains in Lepidoptera

A series of genome assembly and polishing processes were used to construct closed genomes of the *Wolbachia* strains *w*Sca, *w*CauA, *w*CauB, *w*Kue, *w*CI, *w*Ni1, and *w*Hm-c, which were used for the transactivation assays (**Table 2**). Comparative genomics using seven newly sequenced strains and the *w*Hm-t strain (28) identified 788 protein clusters shared by the *Wolbachia* strains, including *Cif*s (*CifB* and *CifA*) and *wmk* homologs (**Figure 2A Table S1**). Proteins that exist in MK *Wolbachia* (*w*Sca, *w*CauA, and *w*Hm-t) but are absent in non-MK *Wolbachia* strains (*w*CauB, *w*Kue, *w*CI, *w*Ni1, and *w*Hm-c) were only the *oscar* homologs encoding ankyrin repeats and papain-like cysteine proteinase domains (**Figure 2A-B**). An *oscar* homolog was also identified in the sequencing data derived from cells coinfected with *w*Fem and *w*CI. Although the genome of the *w*Fem strain was not reconstructed because of technical difficulty in isolating *w*Fem-derived reads from the mixed *w*Fem and *w*CI sequencing data (*w*Fem always coexists with *w*CI and it has not been successfully isolated), *w*CI did not carry *oscar* in its genome (**Figure 2C, Table S1**). Further diagnostic PCR confirmed that the *oscar* homolog was consistently present in wild-caught *E. mandarina* butterflies infected with *w*Fem and *w*CI (*n* = 37) but not in those infected with *w*CI alone (*n* = 77), suggesting that the *oscar* homolog is encoded by *w*Fem. Similarly, *w*Fem-infected females (*n* = 3) of *Eurema hecabe*, a close relative of *E. mandarina*, were *oscar*-positive. Phylogenetic analysis classified the Oscar proteins into two clades: Type I Oscar (Oscar [*w*Fur and *w*Sca] and Ec-Oscar [*w*CauA]) and Type II Oscar (Hm-Oscar [*w*Hm-t] and Em/Eh-Oscar [*w*Fem]; **Figure 2C**). Whereas *Oscar* in the newly sequenced *w*Sca strain was not identical to that in the previously sequenced *w*Sca strain (43), it exhibited high homology to Oscar proteins in the previously sequenced *w*Sca (similarity: 90.4% in 1797 amino acids; bit score: 3241, BLASTp) and *w*Fur (similarity: 94.2% in 1830 amino acids; bit score: 3432) (**Table S1**). Intriguingly, the Type II Oscar homologs Hm-Oscar and Em-Oscar exhibited higher homology than the *w*Sca strains (similarity: 98.5% in 1181 amino acids; bit score: 2362).

**Fig. 2.**
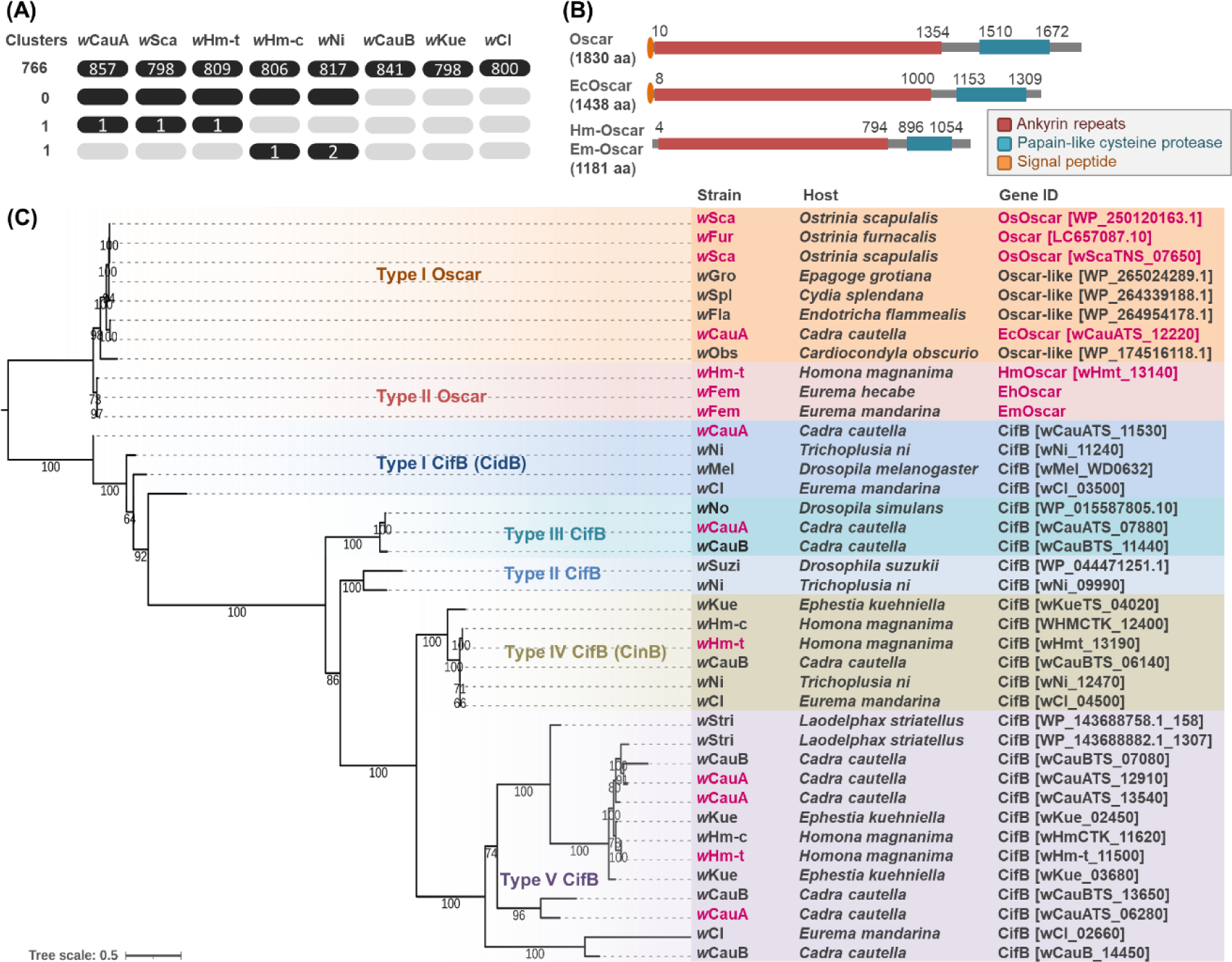
Phylogenies and structures of the Oscar and Cif proteins (A) Protein clusters conserved among the *Wolbachia* strains used in this study. The number of clusters is shown on the left, and the number of proteins contained in the clusters is indicated by black ellipses. (B) Structure of Oscar and its homologs. aa: amino acids. The structure of Oscar (1830 aa), derived from the male killing (MK) *w*Fur strain, is based on Katsuma et al. (26). (C) Phylogeny of Oscar and CifB homologs. Male-killing strains are highlighted in magenta. Accession and gene numbers are given in parentheses.

**Table 2.**
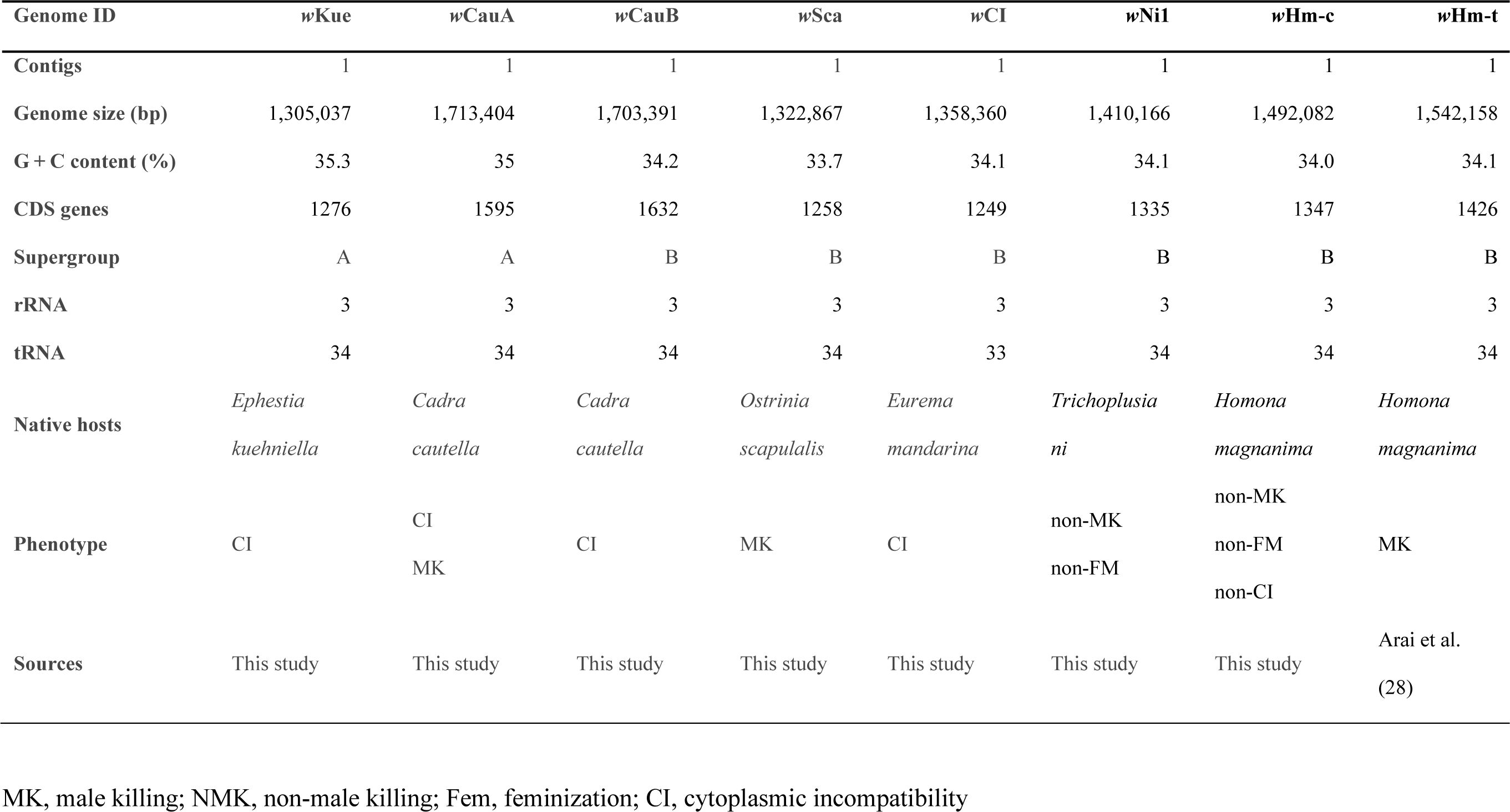
Genomic features of the *Wolbachia* strains used in this study.

### Oscar is absent in non-MK/non-FM *Wolbachia* strains that interact with insect sex determination systems

The *w*Ni1 and *w*Hm-c strains, which induced female-type splicing of *Osmasc* and *Osdsx* in *Ostrinia* male cells, did not carry *oscar* homologs (**Figure 2, Table S1**). In addition, comparative genomics did not identify any proteins that were exclusively present in *Wolbachia* strains that affected the host sex determination system (*w*Hm-t, *w*Sca, *w*CauA, *w*Ni1, and *w*Hm-c) but not in other strains (*w*Kue, *w*CauB, and *w*CI) (**Figure 2A**). A protein cluster specific to *w*Hm-c and *w*Ni1 (i.e., absent in the other *Wolbachia* strains) consisted of three hypothetical proteins (63 amino acids: wHmcTK_11260, wNi_10160, and wNi_12630) that did not encode obvious domains and exhibited no homology to known MK genes (**Tables S1 and S2**). Genes carried by *w*Hm-c or *w*Ni1 that were present in some MK strains (*w*Sca, *w*CauA, and *w*Hm-t) but absent in non-MK strains (*w*CauB, *w*Kue, and *w*CI; **Table S2**) included a *wmk* homolog (wNi_11830), whereas others displayed extremely low or no homology to known MK genes (i.e., Oscar [22], Spaid [44], and PVMKp1 [45]).

## DISCUSSION

Cell transinfection assays demonstrated the inherent feminizing abilities of *Wolbachia* strains. In addition, comparative genomics revealed the presence of *oscar* homologs in these strains, which induced female-specific splicing of three sex-determining genes (*dsx, masc*, and *znf2*) in male *Ostrinia* cells. As previously argued (19, 25), the present study highlighted a common mechanism underlying *Wolbachia*-induced MK and FM in lepidopteran insects, namely Oscar-induced suppression of the *masc,* which results in female-type sex determination and disruption of the dosage compensation system in insects.

Unexpectedly, our study also revealed that two Oscar-deficient *Wolbachia* strains (*w*Ni1 and *w*Hm- c) affected the sex determination system of *Ostrinia*, albeit through a different mechanism from Oscar-bearing *Wolbachia* strains: Oscar-bearing *Wolbachia* strains (i.e., *w*CauA, *w*Sca, *w*Fem, and *w*Hm-t) affected *Osdsx, Osmasc*, and *Osznf2,* whereas Oscar-deficient *w*Ni1 and *w*Hm-c affected *Osdsx* and *Osmasc* but not *Osznf2*. These differences probably arose from the different machineries caused by different *Wolbachia* genes, i.e., *oscar* and other unknown genes. In *B. mori,* Znf2 is an upstream factor in the sex determination cascade that regulates *dsx* expression (24). However, as the sex-determining gene cascade of *Ostrinia* (e.g., the hierarchical relationships among *Osmasc*, *Osznf2*, and *Osdsx*) is not fully understood, it remains unclear why *Osznf2* alone is not affected by *w*Ni1 or *w*Hm-c. One possibility is that Oscar can suppress the functions of *Znf2* (or the cascade containing *Znf2*) as well as *Masc*, whereas the factors carried by *w*Hm-c and *w*Ni1 cannot act on *Znf2* or its upstream components. Our study illustrated that closely related bacteria in the genus *Wolbachia* can activate different functions to manipulate sex determination in a single insect species (**Figure 3**). Further investigation of the causes is warranted.

**Fig. 3.**
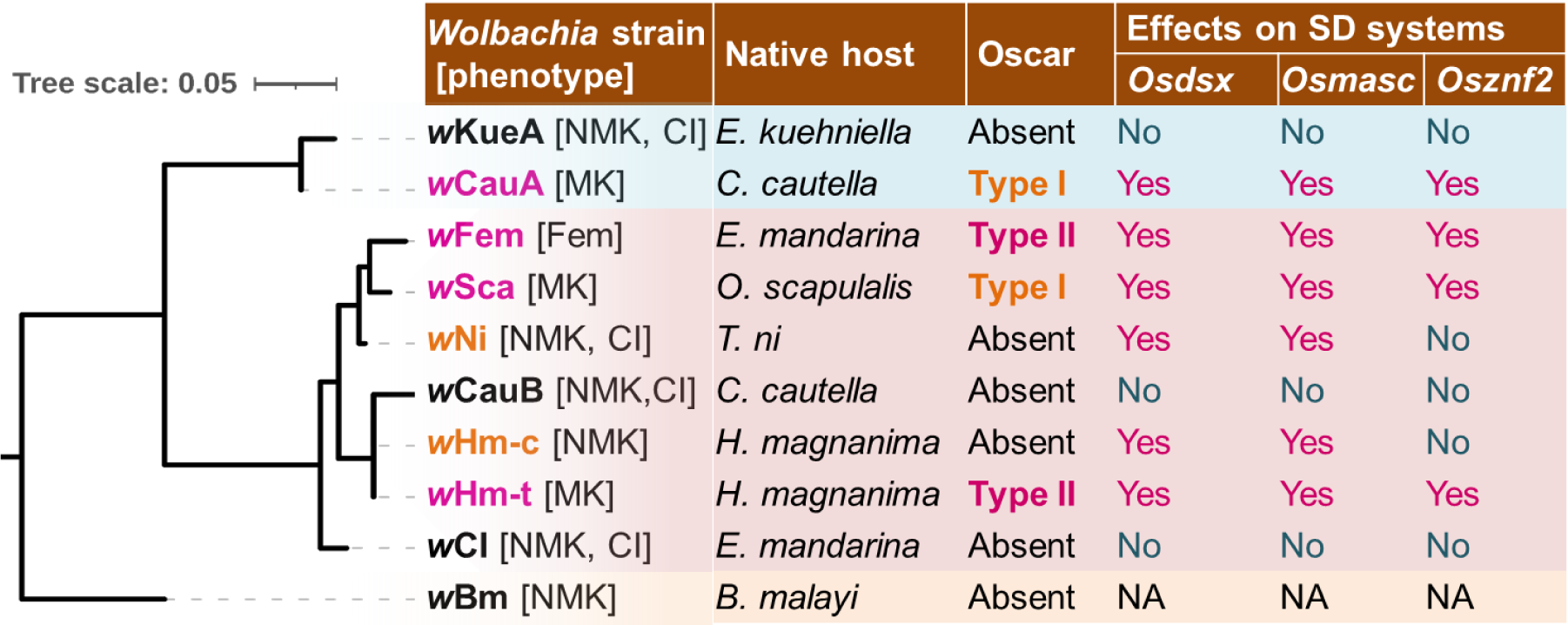
Phylogenies and phenotypes of *Wolbachia* strains and their effects on the sex determination system in *Ostrinia* male cells *wsp* and MLST genes were used to construct the phylogenetic tree. MK/FM strains in native hosts are highlighted in magenta, and strains exhibiting partial feminizing effects are highlighted in orange. The classification of Oscar proteins was based on the phylogeny presented in Figure 2C. The effects on splicing of each sex-determining gene analyzed in this study are presented as either Yes (feminized) or No (not affected). MK, male killing; NMK, non-male killing; Fem, feminization; CI, cytoplasmic incompatibility; NA, not assessed.

This study provided compelling evidence that *Wolbachia* strains (*w*CauA, *w*Ni1, and *w*Hm-c) that do not induce MK or FM in their native hosts (42, 46, Miyata, personal observation) retain the inherent ability to manipulate the sex determination systems of *Ostrinia*. *Wolbachia*-induced reproductive manipulation is influenced by the genetic background of the host (30–33). Under the condition of a female-biased sex ratio, mutations in the host genome that rescue males can be favored by selection. Indeed, the spread of suppressors of symbiont-induced MK has been observed in nature (18, 31, 37–40). For example, the spread of an MK suppressor in the butterfly *Hypolimnas bolina* was almost complete within 5 years (31, 36). Historical records indicate the occurrence of MK (putatively induced by *w*CauA) in the native host *Cadra cautella* until the 1970s (47), but MK does not occur in *w*CauA-infected *C. cautella* (46, 48). However, *w*CauA induces MK when transferred to the closely related host *Ephestia kuehniella* (30). Many species of tortrix moths carry non-MK *w*Hm-c relatives, and phage acquisition is considered responsible for the recent transition from non-MK *w*Hm-c to MK *w*Hm-t in *H. magnanima* (42, 49). The finding that *w*Hm-c retains the ability to influence insect sex determination systems implies that *Wolbachia* have evolved the ability to manipulate insect sex determination systems multiple times. The mechanisms and evolution of MK suppressors should be diverse, as should the mechanism of MK.

In summary, our study demonstrated that the feminizing effect of *Wolbachia* strains on the sex-determining gene cascade is most likely the mechanistic basis of MK and FM in lepidopteran insects. This study highlights the effectiveness of combining cell culture systems and genomic analyses to uncover the inherent ability of *Wolbachia* to manipulate sex. These approaches could also prove effective in elucidating the mechanistic interplay between the host and other selfish reproductive manipulators. A molecular understanding of the commonality and diversity of microbial reproductive manipulations will contribute to a better understanding of the evolutionary interactions between selfish elements and their hosts.

## MATERIALS AND METHODS

### *Wolbachia* strains and insects

*E. kuehniella* lines transinfected with *w*Kue, *w*CauA, and *w*CauB in each were maintained in the laboratory as described by Sasaki et al. (30). Females infected with *w*CauA were mated with uninfected males (48). The insects were reared on a diet consisting of wheat bran, dried yeast, and glycerol (20:1:2) at 25°C under a 16-h/8-h light/dark photoperiod. *H. magnanima* infected with *w*Hm-t (Taoyuan, Taiwan; [41]) or *w*Hm-c (Takao, Japan; [28]) was maintained using Silkmate 2S (Nosan Co., Yokohama, Japan). The MK host line W^T12^ was maintained by crossing it with a *Wolbachia*-free normal sex ratio line, as described by Arai et al. (41). For *E. mandarina,* all-female blood coinfected with *w*Fem and *w*CI and a normal sex ratio line singly infected with *w*CI were collected on Tanegashima Island, Japan and maintained on an artificial diet, as reported by Kageyama et al. (25). *T. ni* harboring *w*Ni1 was collected in Matsudo, Chiba, Japan. The MK *w*Sca strain, maintained in BmM2 cells as described by Herran et al. (23), was used for the following assays.

### Transinfection of *Wolbachia* into cultured cells

Fat bodies septically isolated from *Wolbachia*-infected insects were placed in the cell lines AeAl2 (*w*CauA, *w*CauB, *w*Kue, *w*Ni1, *w*Hm-t, and *w*Hm-c) and BmN4 (*w*CI and *w*Fem). After confirming that *Wolbachia* was stably maintained by diagnostic PCR, *Wolbachia*-positive AeAl2/BmN4 cells were used as donors for *Wolbachia* transinfection into *the O. scapulalis* M1 (OsM1) cell line. *Wolbachia*-infected AeAl2/BmN2 cells were passed through a 5.0-µm filter (Cat. No. FJ25ASCCA050PL01, GVS, Via Roma, Italy), and six drops (approximately 300 µL) of the flow- through were added to the recipient OsM1 cell line.

### RT-qPCR

Total RNA (200–500 ng) extracted from harvested cells using Isogen II (Nippon Gene, Tokyo, Japan) or TRIzol RNA Isolation Reagent (Thermo Fisher Scientific, Waltham, MA, USA) was reverse-transcribed using a PrimeScript II 1st strand cDNA Synthesis Kit (Takara Bio, Shiga, Japan) according to the manufacturer’s protocol. qPCR was performed with 5 µL of KOD SYBR (Toyobo), 0.4 µL each of the forward and reverse primers (10 pmol/µL), 2.2 µL of water, and 2.0 µL of the cDNA template. qPCR was performed in a LightCycler 96 System (Roche, Basel, Switzerland) with a temperature profile of 180 s at 95°C; 40 cycles of 8 s at 98°C, 10 s at 60°C, and 10 s at 68°; and heating to 90°C for melting curve analysis. The primers used in this study are listed in **Table S3**. The relative expression (*Osmasc*, *Osdsx*, and *Osznf2* versus the control gene *Osef1a*) and the ratio of male-to-female splice variants were estimated.

### Genomic analyses

*w*Sca (in BmM2 cells, described by Herran et al. [23]), *w*Hm-c, *w*Ni1, *w*CauA, *w*Kue, *w*CI, and *w*Fem and *w*CI (i.e., coinfected into AeAl2 cells) were purified from cells harvested in 150-mL flasks as described by Iturbe-Ormaetxe et al. (50) with several modifications. Cells were pelleted by centrifugation (900 × *g* for 5 min) and homogenized with glass beads using Multi-beads Shocker MB3000 at 2000 rpm for 40 s (Yasui Kikai Co. Osaka, Japan). The lysates were passed through 5.0- and 1.2-µm filters and centrifuged at 15,000 × *g* for 30 min at 4°C. Pellets containing *Wolbachia* cells were subjected to DNA extraction using a NanoBind Big DNA Tissue kit (Circulomics, Baltimore, MD, USA) according to the manufacturer’s protocol. The extracted DNA was sequenced using an Illumina short read (paired-end 150 bp) and MinION with a Rapid sequencing kit (Oxford Nanopore, Oxford, UK) with a Nanopore flange flow cell (R9.4) (Oxford Nanopore). For *w*CauB, DNA extracted from host insects harboring *Wolbachia* using a NanoBind Big DNA Tissue kit was sequenced using Illumina (paired-end 150 bp) and Nanopore platforms (ligation sequencing kit with Nanopore MinION flow cell [R10.4], Oxford Nanopore). The Nanopore data were assembled using Flye v1.6 (51) and polished 3–5 times with Illumina data using Pilon (52) and minimap2 (53). The resulting circular *Wolbachia* genomes were annotated using DFAST (54). Protein homologies were analyzed using OrthoVenn 3 (https://orthovenn3.bioinfotoolkits.net). *Wolbachia cifB*, *wmk*, *and oscar* (accession numbers are presented in **Figure 2** and **Table S1**) (26, 29, 55) were used as queries to identify homologs from *Wolbachia* genomes using local BLASTn and BLASTp searches (default parameters). Motifs in *oscar* homologs were surveyed using InterPro (https://www.ebi.ac.uk/interpro/) and HHpred (https://toolkit.tuebingen.mpg.de/tools/hhpred). *cifB* and *oscar* were aligned using ClustalW (56), trimmed using trimAl (57) in the default mode, and used to construct a phylogenetic tree based on maximum likelihood with bootstrap resampling of 1000 replicates using the IQTREE server (http://iqtree.cibiv.univie.ac.at/).

### PCR detection of *Em/Eh-oscar* gene

*E. mandarina* collected from Tanegashima Island (Kagoshima, Japan) (Kageyama et al., 2020) and *E. hecabe* collected from Kohama Island (Okinawa, Japan) were subjected to DNA extraction using a DNeasy kit (Qiagen, Hilden, Germany) according to the manufacturer’s protocol. The DNA concentration was adjusted to 10 ng/µL and subjected to PCR using primer sets amplifying the complete sequences of *Em/Eh-oscar* and *Hm-oscar* (**Table S3**). *Em/Eh-oscar* was amplified using Emerald Amp Max Master Mix (TaKaRa) at 94°C for 3 min; 35 cycles of 94°C for 30 s, 62°C for 30 s, and 72°C for 3 min; and a final extension at 72°C for 7 min, as described by Arai et al. (28).

### Statistical analysis

For RT-qPCR assays, we used the average cycle threshold value (Ct) for each sample and estimated the relative expression, as described by Herran et al. (23) and Sugiomoto et al. (20). Elongation factor a (Ef1a) was used as the control gene. The 2^−ΔCt^ (i.e., CtAve target gene − CtAve ef1a) and 2^−ΔCtMal^2^−ΔCtFem^ (i.e., relative expression in males vs. females) values were calculated. The expression ratios of the splice variants (i.e., *DsxM/DsxF*, *MascM/MascF*, and *Znf2M/Znf2F*) were analyzed using the Steel–Dwass test in R software v4.0 (58).

### Data accessibility

The sequence read data are publicly available in DDBJ under the accession numbers PRJDB16497 (BioProject), DRA016973 (DDBJ Sequence Read Archive: DRA), and DRA017337 (DRA). *Wolbachia* genomes (contigs) are available in the DDBJ database under the following accession numbers: *w*CauA (SAMD00639214, AP028948), *w*CauB (SAMD00639215, AP028949), *w*Kue (SAMD00639216, AP028950), *w*CI (SAMD00639217, AP028951), *w*CI-Fem (SAMD00639218, AP028952), *w*Sca (SAMD00639219, AP028953), *w*Ni1 (SAMD00639220, AP028954), and *w*Hm-c (SAMD00654377, AP028994).

## Acknowledgements

We acknowledge the support from the Japan Society for the Promotion of Science (JSPS) Research Fellowships for Young Scientists [Grant Number 20F40719, 21J00895], JSPS Grant-in-Aid for Scientific Research [Grant Number 20K06084, 22K14902, 22K20663, 23H02229], Cabinet Office, Government of Japan, Moonshot Research and Development Program for Agriculture, Forestry, and Fisheries (funding agency: Bio-oriented Technology Research Advancement Institution) (no. JPJ009237).

## Author Contributions

H.A. collected and maintained the insects, transfected Wolbachia, conducted genome sequencing, qPCR, and data analysis, wrote the original manuscript, and revised the manuscript. B.H. transfected Wolbachia, conducted qPCR, and contributed to the discussion. T.N.S. collected insects, transfected *Wolbachia*, conducted qPCR, and contributed to the discussion. M.M. collected insects and contributed to the discussion. T.S. collected and maintained the insects and contributed to the discussion. D.K. transinfected *Wolbachia*, designed and conceptualized experiments, and revised the original manuscript.

## Inclusion and diversity

We support inclusive, diverse, and equitable research.

## Legends of the supplementary material

Table S1 Homologies between phenotype-associated *Wolbachia* proteins and proteins of the *Wolbachia* strains used in this study based on BLASTp searches.

Table S2 *w*Hm-c- or wNi1-encoded strain-specific proteins and proteins shared with any of the MK *Wolbachia* strains.

Table S3 Primers used in this study.

